# Context-specific modulation of intrinsic coupling modes shapes multisensory processing

**DOI:** 10.1101/509943

**Authors:** Edgar E. Galindo-Leon, Iain Stitt, Florian Pieper, Thomas Stieglitz, Gerhard Engler, Andreas K. Engel

## Abstract

Intrinsically generated patterns of coupled neuronal activity are associated with the dynamics of specific brain states. Sensory inputs are extrinsic factors that can perturb these intrinsic coupling modes, creating a complex scenario in which forthcoming stimuli are processed. Studying this intrinsic-extrinsic interplay is necessary to better understand perceptual integration and selection. Here, we show that this interplay leads to a reconfiguration of functional cortical connectivity that acts as a mechanism to facilitate stimulus processing. Using audiovisual stimulation in anesthetized ferrets, we found that this reconfiguration of coupling modes is context-specific, depending on long-term modulation by repetitive sensory inputs. These reconfigured coupling modes, in turn, lead to changes in latencies and power of local field potential responses that support multisensory integration. Our study demonstrates that this interplay extends across multiple time scales and involves different types of intrinsic coupling. These results suggest a novel large-scale mechanism that facilitates multisensory integration.

## INTRODUCTION

Large-scale patterns of intrinsically generated coupling are a hallmark of brain networks. These coupling patterns, which we term intrinsic coupling modes (ICMs) (1), have gained attention since they may have crucial importance for brain function (1–9). ICMs correspond to dynamic coupling patterns which are not imposed by entrainment to an external stimulus or movement but emerge from the connectivity of cortical and subcortical networks (1). Substantial evidence suggests two different mechanisms for ICMs. One type (which we term phase ICMs) arises from phase coupling of band-limited oscillatory signals, whereas the other results from coupled aperiodic fluctuations of signal envelopes (hence termed envelope ICMs) (1). Importantly, ICMs occur both in ongoing activity and during tasks. Their impact has been observed in the modulation of sensory responses (10–13), behavior and perception (14–18) and, furthermore, ICMs play a key role in mediating effective and selective communication in networks of cortical and subcortical areas (19–22).

Cortical activity is believed to arise from the complex interplay between the intrinsic dynamics and the continuous stream of sensory information impinging from the external environment (23, 24). Understanding this interplay is fundamental for elucidating the role of ICMs in information processing. In a classical approach the global organization of neural activity, or brain state, has been viewed as the arena or playground on which extrinsic sensory inputs are processed (2, 24–27). The opposite, that is, the effects of sensory stimuli on ongoing activity is less well understood. Whereas Fiser et al. (5) found that during visual stimulation most of the spatiotemporal correlations in visual cortex are driven by the network and not by the stimulus, a study by He using fMRI (28) revealed a significant negative interaction between ongoing and evoked activity. Thus, the large-scale interaction between intrinsic and externally driven dynamics is still far from being understood.

By using a 64-channel electrocorticographic (ECoG) array, we study here the interplay between large-scale ICMs and extrinsically evoked activity during repeated multisensory stimulation in the anesthetized ferret. To elucidate this interplay, two complementary aspects were analyzed. On the one hand, we investigated how visual, auditory and audiovisual stimuli modulated ICMs. On the other hand, we tested whether phase or envelope ICMs, extracted from pre-stimulus intervals, predicted stimulus-related multisensory effects. We found that pre-stimulus functional connectivity affects both response timing and power, and that this effect extends over several frequency bands of cortical activity. Interestingly, this causal correlation is itself stimulus-context dependent, suggesting an adaptation in functional connectivity giving rise to ICMs. Our results suggest that the role of this connectivity adaptation is to facilitate multisensory integration in the cortex.

## RESULTS

General procedures related to surgical preparation and data acquisition have been described in detail elsewhere (22). Briefly, we recorded cortical activity in ferrets using custom-built ECoG arrays (29) with 64 electrodes distributed to cover the posterior cortex of the left hemisphere in the ferret (Fig. 1A,B; Fig. S1A,B). For the present study a total of 5 adult female anesthetized ferrets (Mustela putorius) were used (see Methods). We collected data during stimulus presentation (see Methods) and from periods of ongoing activity. For the stimulus-related study we used two classes of stimulus blocks: transient stimuli (clicks and flashes) and sustained complex stimuli (drifting Gabor patches and auditory ripples). This allowed us to determine whether possible changes in ICMs are stimulus-specific. For statistical analysis we grouped the responsive sites into auditory areas (A1, AAF, ADF, AVF, PSF and PPF) and visual areas (17, 18, 19, 20, 21 and SSY) (Fig. 1A).

**Figure 1.**
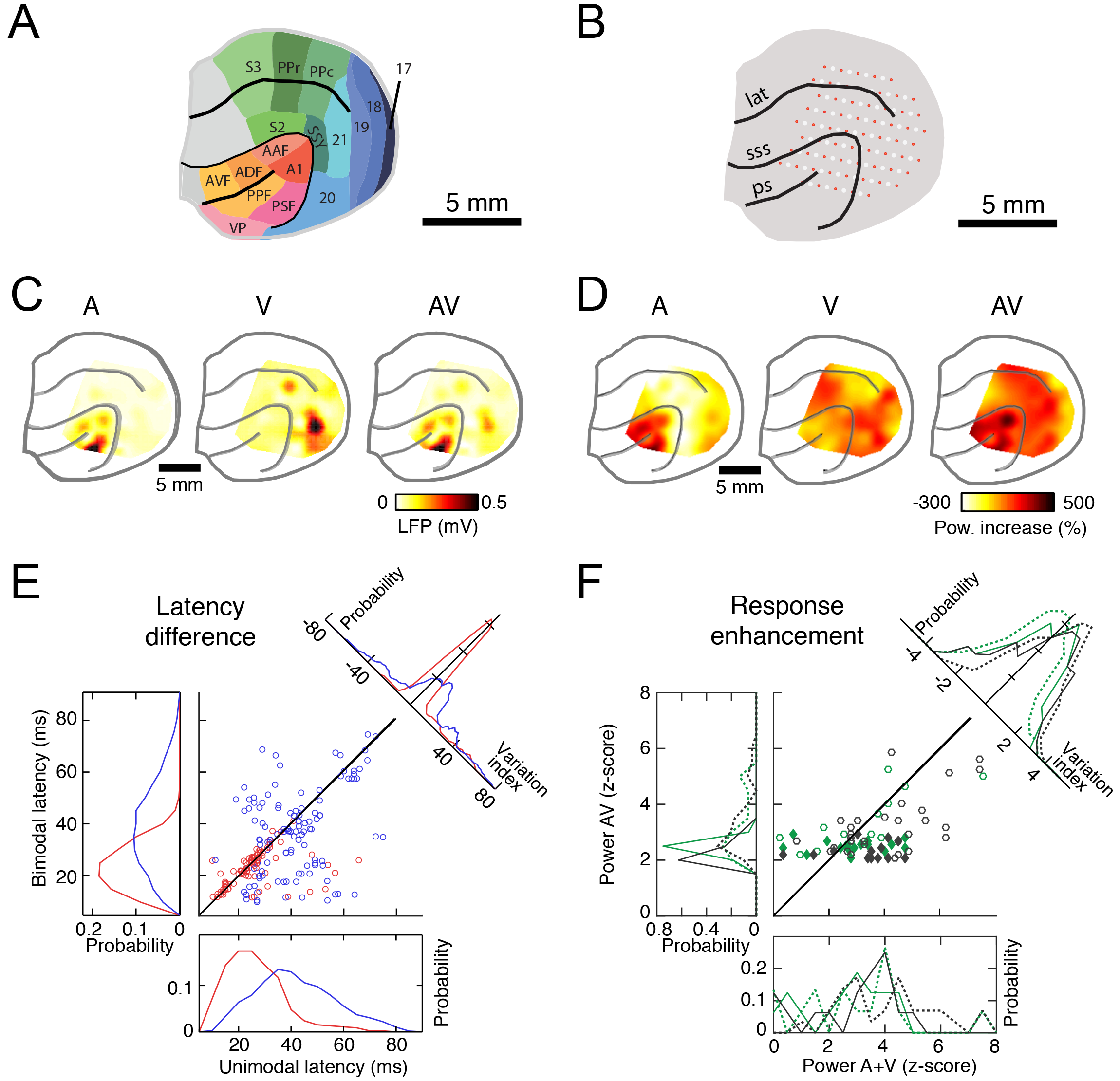
Effects of multisensory stimulation on response timing and power. (**A**) Map of functional areas. Abbreviations: A1, primary auditory cortex; AAF, anterior auditory field; PPF, posterior pseudosylvian field; PSF, posterior suprasylvian field; ADF, anterior dorsal field; AVF, anterior ventral field; VP, ventroposterior area; S2, primary somatosensory cortex; S2, secondary somatosensory cortex; S3, tertiary somatosensory cortex; PPc, posterior parietal caudal; PPr, posterior parietal rostral; SSY, suprasylvian visual areas. (**B**) Schematic of the ECoG array placed on the left hemisphere of the ferret brain, covering most of the occipital, temporal and parietal cortex. Red dots represent recording sites, and white circles represent the holes in the foil. Abbreviations: lat, lateral sulcus; sss, suprasylvian sulcus; ps, pseudosylvian sulcus. (**C**) Topographic distribution of response amplitudes with unimodal (A: clicks, V: flashes) and bimodal (AV: simultaneous click and flashes) stimuli. Dark colors represent the amplitude of the strongest deflection in the event-related potential (ERP) within the first 80ms after stimulus onset. (**D**) Topographic distribution of changes of total power in the alpha-band in response to auditory (left), visual (middle) and audiovisual (right) stimuli. Power change was averaged over a stimulus time window (100 to 600 ms, relative to stimulus onset) and normalized to the pre-stimulus interval (−600 to −100 ms). (**E**) Latency reduction. Scatter plot of latencies of responses to unimodal versus bimodal stimulation. Blue circles represent data from electrodes that responded to unimodal visual stimuli, but not unimodal auditory stimuli; red circles indicate electrodes that responded in the unimodal condition only to auditory stimuli. Sites that responded to both unimodal visual and unimodal auditory stimuli were not included. Top right: probability distribution of the variation index (difference between latencies with uni- and bimodal stimuli). Elements with low variation index are located along the diagonal. Bottom: probability distribution of unimodal latencies. Left: probability distribution of bimodal latencies. (**F**) Scatter plot of power enhancement (z-score) during unimodal (ripples or drifting Gabor patches) and bimodal stimulation across responsive electrodes for theta band (4-8 Hz, green circles) and alpha band (8-16 Hz, black circles) for sustained stimuli. Continuous lines and filled diamonds correspond to visual areas, while dashed lines and open circles correspond to auditory areas. Top right, normalized probability distribution of the variation index (difference between unimodal and bimodal power enhancements). Bottom and left, panels showing probability distributions of power enhancement during unimodal and bimodal stimuli, respectively.

### Multisensory interactions modulate response latency and power

First, in order to study the role of ICMs in stimulus processing it was necessary to characterize the spatial distribution and the multisensory properties of cortical sensory responses in the ferret. Figure 1C shows the spatial distribution of amplitudes of event-related potentials (ERPs) to auditory, visual and combined audiovisual stimulation in a single animal. Across animals, the topography of cortical responses was in agreement with previous studies on auditory (30, 31) and visual stimulation in the ferret (32). A total of 92 recorded sites responded to auditory stimuli across animals, while 118 sites responded to visual stimuli. To test for multisensory effects on response timing we compared the ERP latencies between uni- and bimodal stimuli (see Methods). Visual responses showed higher latency variability compared to auditory responses (Avar = 139 ms and Vvar = 45 ms, Brown-Forsythe test, p < 0.0001). Most importantly, bimodal stimulation resulted in a significant latency reduction for visually responsive sites (mean V = 40.2 ms, AV = 34.3 ms; sign test, *p* = 1.7×10-5) but not for sites with auditory responses (A = 24.6 ms, AV = 20.3 ms; sign test, *p* = 0.44). This effect was particularly pronounced for sites along the suprasylvian sulcus and a region at the pseudosylvian sulcus, which responded to visual stimuli, and has been suggested as the homologue of the cat’s anterior ectosylvian visual area (AEV) located in the multisensory ectosylvian sulcus (AES) (31, 33, 34).

In addition, we analyzed the spectral distribution of stimulus-related changes in normalized total power (see Methods). As shown by mean time-frequency spectrograms (Fig. S2B), stimulus-related changes in total power were observed across a broad range of frequencies with all stimuli employed. Figure 1D illustrates the topography of cortical stimulus-related total power in the alpha band (8-16 Hz) in a single animal. Generally, although the spatial distribution of responses agreed well with known functional properties of cortical areas, the number of responsive sites in auditory areas varied across frequency bands (Fig. S3 and S4).

Multisensory effects in LFP power responses were quantified by the difference AV-(A+V), with A, V and AV being the LFP normalized total power in response to auditory, visual and audiovisual stimulation, respectively. To test the effects of sensory stimulation we used blocks with either sustained stimuli (ripples and/or gratings, R-G) or short transient stimuli (clicks and/or flashes, C-F; for details, see Methods). In transient stimulation blocks, multisensory effects in auditory areas were frequency-dependent (one-way ANOVA, F _1,130_ = 6.5; *p* < 0.001 with Bonferroni correction for frequency) and mostly dominated by subadditive effects across frequency bands (Fig. S3E). Super-additive effects occurred more rarely at both low and high-frequencies. Similarly, in visual areas the subadditive effect dominated across both stimulus blocks and frequency bands (Fig. S3F). In sustained stimulation blocks, multisensory effects were frequency dependent (Fig. 1F; Fig. S3G,H); in particular, the difference in power (AV-A-V) in alpha, beta, and gamma band was significantly lower than zero (t-test, *p* < 0.001, with Bonferroni correction). The population distribution of multisensory responses and the topography of the multisensory effects across all frequency bands are shown in Figures S3 and S4. Altogether, our results show that multisensory effects on response power are dominated by subaddition, with less sites showing superadditivity. Similar observations have been made by Kayser et al. (35) in auditory cortex of monkeys. Moreover, our data show that the multisensory effects on the power of responses differ between the two types of stimulus blocks for both, auditory and visual areas (one-way ANOVA F_1,334_ = 6.6, *p* = 0.01 and F_1,600_ = 17.6, *p* < 0.0001, respectively).

### Sensory stimuli have limited impact on ICMs

A key goal of our work was to study whether sensory stimulation modulates ICMs in the cortex. We tested this first for envelope ICMs which were quantified by correlation of the signal amplitudes. For computing this coupling measure, we use the orthogonalized components of the respective signals to eliminate amplitude envelope correlations introduced by volume conduction (36).

We computed connectivity matrices for periods between 100 and 600 ms after stimulus onset for the stimulation blocks with sustained and those with transient stimulation. Electrodes in the array were reordered and grouped according to their modular structure in order to build the connectivity matrix (Fig. 2). Matrices in Figure 2 show the patterns of connectivity during the presentation of auditory, visual and audiovisual stimulation. To evaluate whether these correlations represent functional connectivity, or rather artifactual correlations through common inputs, we compared these matrices with the connectivity obtained after shuffling trials within each stimulus modality. Connections that were not significantly different from these shuffle controls are marked with a cross (x) in the coupling matrices in Fig. 2B and Fig. 3A,B.

**Figure 2.**
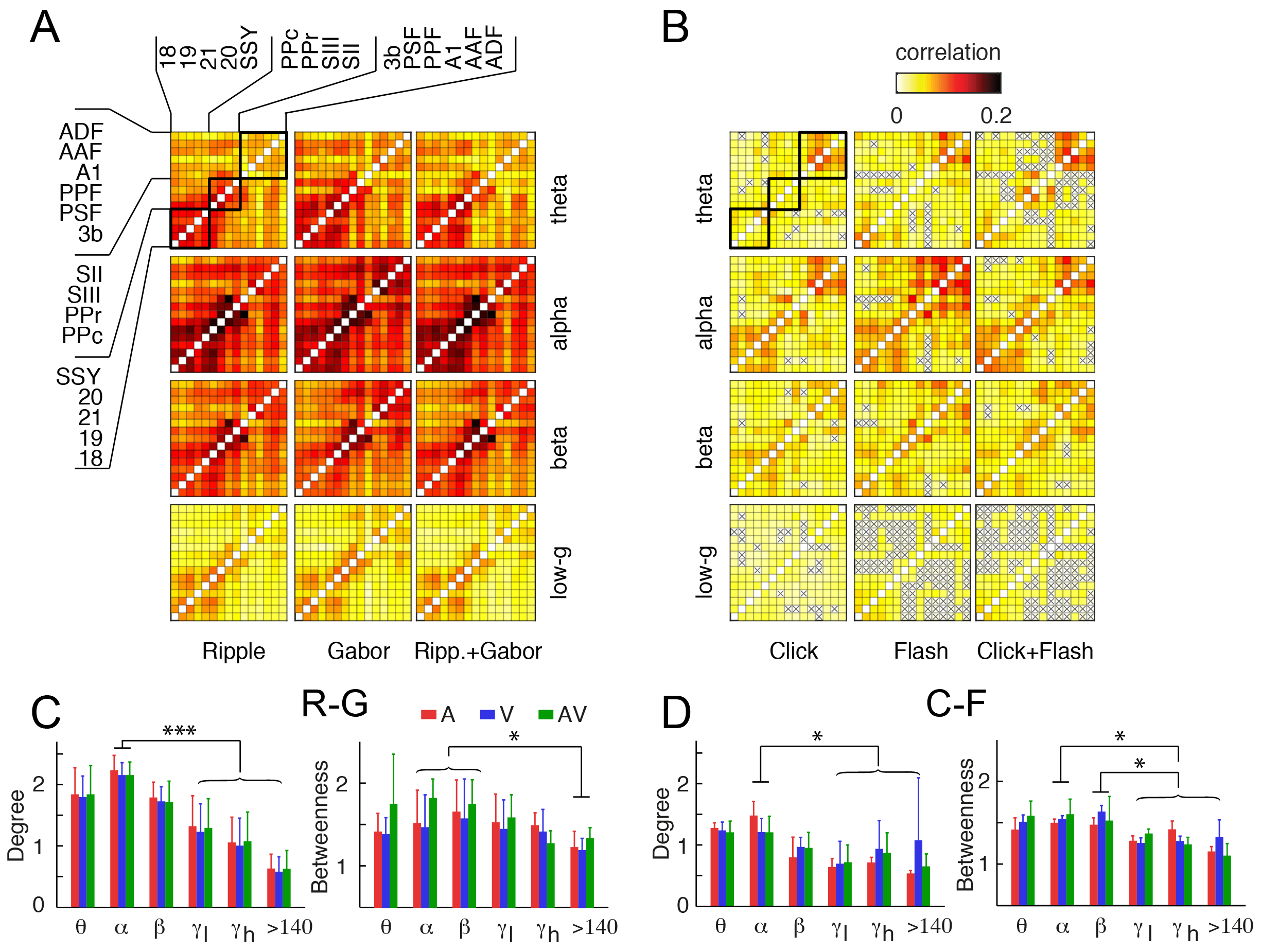
Stimulus-related modulation of envelope ICMs. Coupling matrices represent the strength of amplitude envelope correlations between functional areas. The labeling of rows and columns, as shown in the top left in panel (**A**), applies to all matrices in the figure. Based on the anatomical regions (abbreviations as in Fig. 1), these can be grouped in three modalities: visual, somatosensory-parietal and auditory regions (black dark rectangles in top-left matrix). (**A**) Connectivity during stimulation with ripples (left column, A), drifting Gabor patches (second column, V), and simultaneous presentation of ripples and Gabor patches (third column, AV). Different rows represent different frequency bands. Note that matrices for the high-gamma and the high-frequency bands are not shown. (**B**) Connectivity during unimodal (A, V) and bimodal (AV) stimulation with clicks and flashes for different frequency bands (rows). Connections that were not significantly different from the shuffle controls are marked with an “x”. (**C**) Normalized degree (left) and betweenness (right) for connectivity matrices shown in (**A**). Each subplot represents the mean and standard error of the respective graph measure associated with auditory (red), visual (blue) and audiovisual (green) stimulation. (**D**) Normalized degree and betweenness for connectivity matrices shown in (**B**).

**Figure 3.**
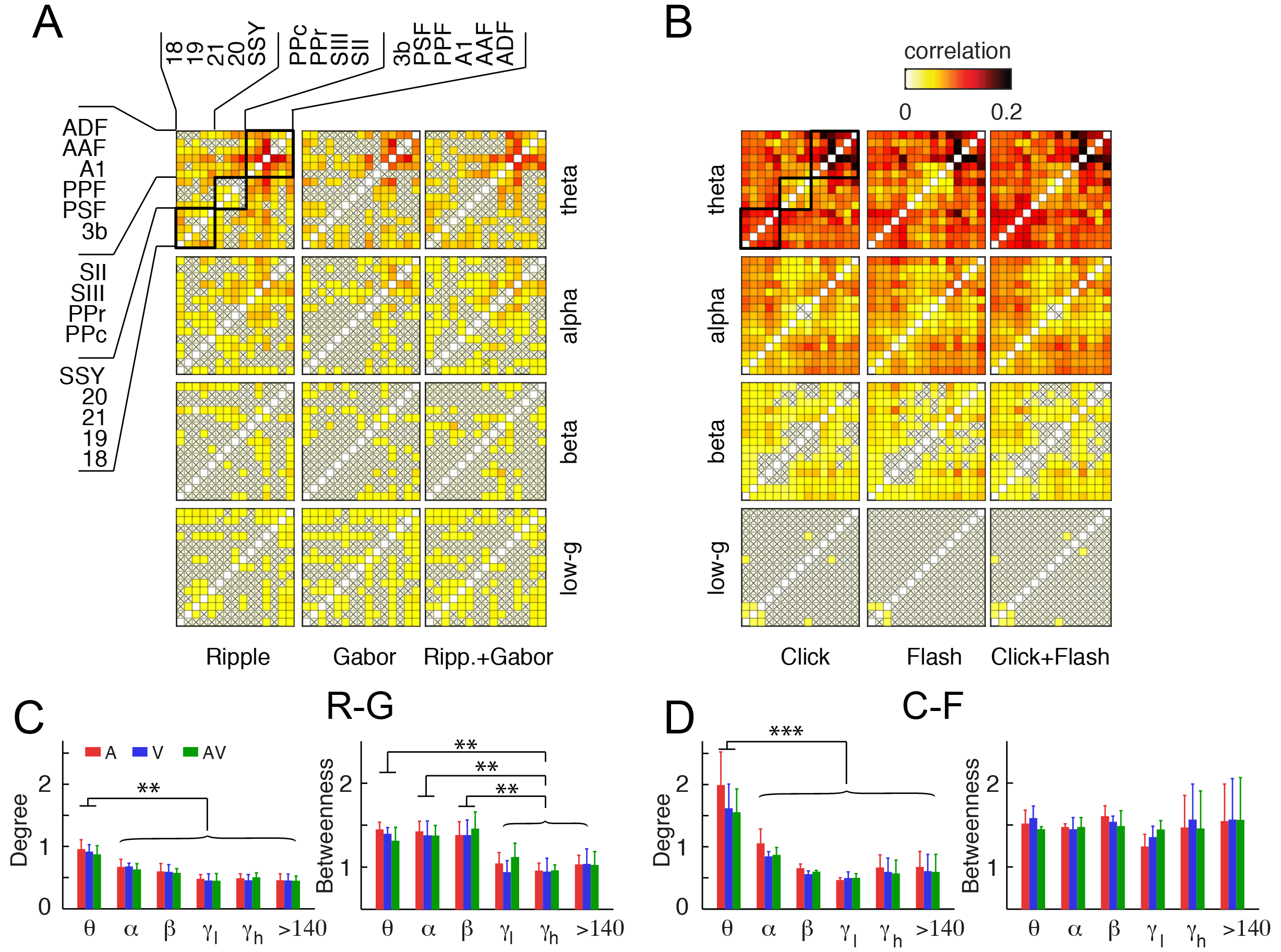
Stimulus-related modulation of phase ICMs. (**A**) Matrices from the first to the third column represent the mean imaginary coherence between areas during stimulation with auditory ripples (A) and drifting Gabors (V), or both (AV). (**B**) Connectivity during unimodal (A, V) and bimodal (AV) stimulation with clicks and flashes. In (A) and (B), connections that were not significantly different from the shuffle controls are marked with an “x” in the coupling matrices. (**C**) Normalized degree and betweenness connectivity matrices shown in (**A**). (**D**) Normalized degree and betweenness for connectivity matrices shown in (**B**).

Strongest correlations appeared mostly within clusters of functionally related areas, rather than across modalities. Furthermore, within blocks the connectivity did not differ significantly between stimulation conditions (Fig. 2A and 2B). In general, we observed higher envelope ICMs for low frequency (theta, alpha, beta) compared to high frequency bands. Importantly, within the sustained block the connectivity matrices did not differ significantly, on an area-by-area basis, from those in the pre-stimulation intervals (Mann-Whitney U test at a two-sided *p* < 0.05, Bonferroni corrected) (Fig. 2A, cf. Fig. 4A). In this stimulation block, all connections were significantly stronger than the shuffled condition.

**Figure 4.**
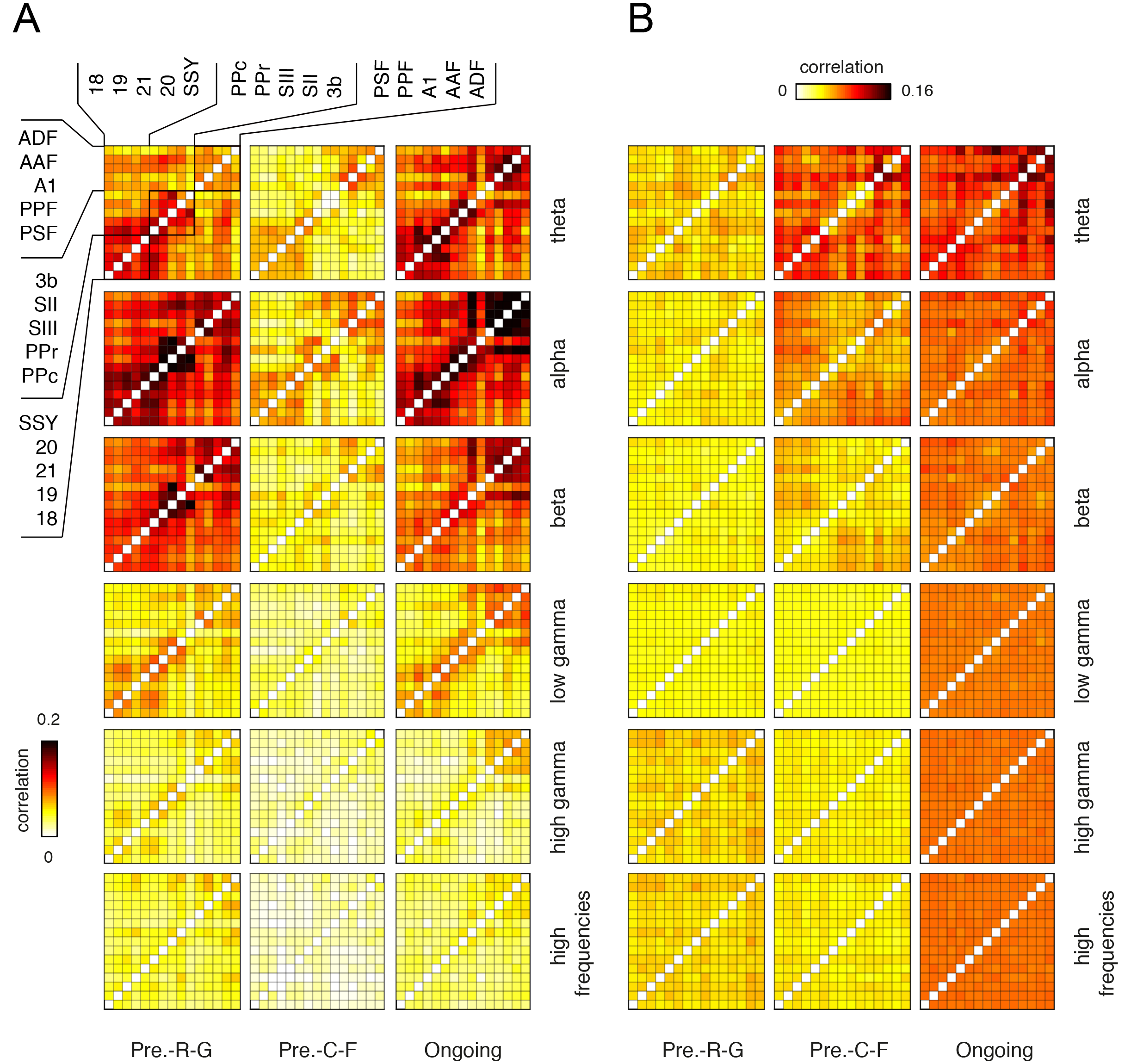
Pre-stimulus connectivity matrices. (**A**) Envelope ICMs. Left column shows the average matrix across animals of the pre-stimulus connectivity obtained during the sustained stimulation blocks (pre-RG). Middle column shows the pre-stimulus connectivity matrix for the blocks with clicks and flashes (pre-CF). Right column: connectivity in the ongoing condition, i.e, in a recording block without intermittent sensory stimulation. Rows represent the frequency bands as defined in Methods. (**B**) Phase ICMs. Matrices show imaginary coherence in pre-RG, pre-CF and ongoing activity blocks for the same recordings epochs as in (**A**).

We used graph-theoretical measures to investigate whether sensory stimulation alters the topology of ICMs. In principle this could take place in different ways, for example, through a global modulation in strength of connectivity or through topological reorganization of the functional network architecture. To quantify these effects we computed average connectivity strength (degree), prevalence of hubs (betweenness) and network segregation (clustering coefficient) (37). Strength of connectivity was not affected by stimulus modality (two-way ANOVA, *p* = 0.93), however, it did show a dependency across frequency bands (two-way ANOVA, F_1,189_ = 22.2, *p* = 0.04, Bonferroni corrected for frequency). The strongest correlations were observed in the alpha-band, where degree was significantly higher than in the gamma-bands (*p* = 1.3×10^−6^ and *p* = 2.07×10^−7^, t-test, Bonferroni corrected) and high-frequency band (t-test, *p* = 1.2×10^−8^, Bonferroni corrected). In contrast, betweenness was neither affected by stimulus modality nor frequency bands (two-way ANOVA, *p* > 0.3, in both cases) (Fig. 2C). Finally, stimulus block showed no effect in the network segregation across frequency bands (Fig. S5).

Applying the same analysis to transient-stimulation blocks we observed, again, that connectivity was not influenced by sensory modality (Fig. 2B). As in the sustained block, coupling was higher within functional cortical systems. In the transient-stimulation blocks, some connections were not significantly stronger than the shuffled condition, especially at high frequencies (Fig. 2B). However, such connections were rather weak, suggesting that functional coupling generally was not introduced through common inputs. Both graph measures in Fig. 2D did not appear to reflect the stimulus modality (two-way ANOVA, F_1,107_ = 0.11, *p* = 0.89). Here, again, average connectivity strength was frequency band specific (two-way ANOVA, F_1,107_ = 6.9, *p* < 0.01, Bonferroni corrected). Finally, a comparison between stimulus blocks showed a significant effect on connectivity strength (one-way ANOVA, F_1,197_ = 23.2, *p* < 10^−5^, Bonferroni corrected), as well as on clustering properties (Fig. S5).

In the same dataset, phase ICMs were quantified by using the imaginary part of the coherence (38) (Fig. 3). This analysis revealed connectivity patterns that strongly differed from those obtained by envelope correlation. Strongest phase ICMs were found mainly in the theta and alpha band. They were most prominent among auditory areas and, in the C-F blocks, also between visual and auditory regions. Importantly, as for envelope ICMs, the impact of stimulation modality on phase ICMs was not significant neither in R-G nor C-F blocks (Fig. 3C and 3D). In contrast connectivity strength, betweenness and clustering coefficient appeared to be suitable to detect differences between stimulation blocks (two-way ANOVA, F_1,214_ = 21.4, *p* < 0.001 and F_1,214_ = 17.1, *p* < 0.001, respectively). Furthermore, clustering coefficients (Fig. S5) significantly differed between frequency bands (two-way ANOVA, *p* < 10^−5^; ANOVA) but not between stimulus modalities. As shown by cross (x) labelling in Fig. 3A and B, we also observed connections for which phase coupling did not differ significantly from shuffle controls. However, this only held for connections with very low values of imaginary coherence.

Taken together, our data consistently show that during sensory stimulation, significant functional connectivity patterns can be observed that exhibit spatial specificity and differ across frequency bands. Interestingly, however, these coupling patterns do not reflect the modality of the presented stimulus. A possible explanation for this unexpected result might be that the types of ICMs investigated here mostly reflect the underlying structural connectivity of the networks under study. Alternatively, they might be shaped by factors that influence functional connectivity on a longer time scale, such as changes in neuromodulatory systems, more slowly occurring state changes in the network, or contextual effects resulting from repeated sensory stimulation over extended time periods. In the latter case, a relevant parameter in our dataset was the stimulus block, i.e., the repeated exposure to sustained or transient stimuli.

### Pre-stimulus ICMs are context specific

We therefore decided to test whether the connectivity patterns during intervals with no stimulation were dependent on the stimulus block they were embedded in. We hypothesized that, if these differed between blocks, this would support the idea of modulation of connectivity through contextual effects. The time interval was the same duration as before (500 ms) but now taken from −600 ms to −100 ms relative to stimulus onset. We analyzed connectivity matrices for pre-stimulus intervals across stimulation blocks and frequency bands using the same connectivity measures as used before (Fig. 4). Generally, we observed strong differences in the spatial patterning of envelope and phase ICMs. In most frequency bands, envelope ICMs showed a clustering of areas into functional systems. This was particularly pronounced, e.g., for the pre-stimulus intervals of sustained stimuli in the alpha band (Fig. 4A). Interestingly, for envelope ICMs, there were strong differences between the mean connectivity strengths of the two stimulus blocks, especially at low frequencies (t-test, *p* < 10^−8^ for all cases, Bonferroni corrected). In contrast, for phase ICMs the mean strength in pre-stimulus intervals of the transient stimulation blocks were stronger for theta, alpha, and beta bands (t-test, *p* < 10^−5^ in all cases) (Fig. 4B). However, for gamma bands and the high frequency band, coherence was higher preceding sustained stimuli than transient stimuli (t-test, *p* < 10^−5^). For both envelope and phase ICMs, rescaling of the connectivity matrices by normalizing to the mean connectivity (Fig. S6) emphasized the similarity of coupling patterns across frequency bands within each of the two ICM classes, suggesting that part of the difference emerges from a modulation of connectivity strengths. It should be noted that graph-theoretical analysis for the pre-stimulus intervals generally yielded the same results as that for the stimulation epochs (Fig. 5).

**Figure 5.**
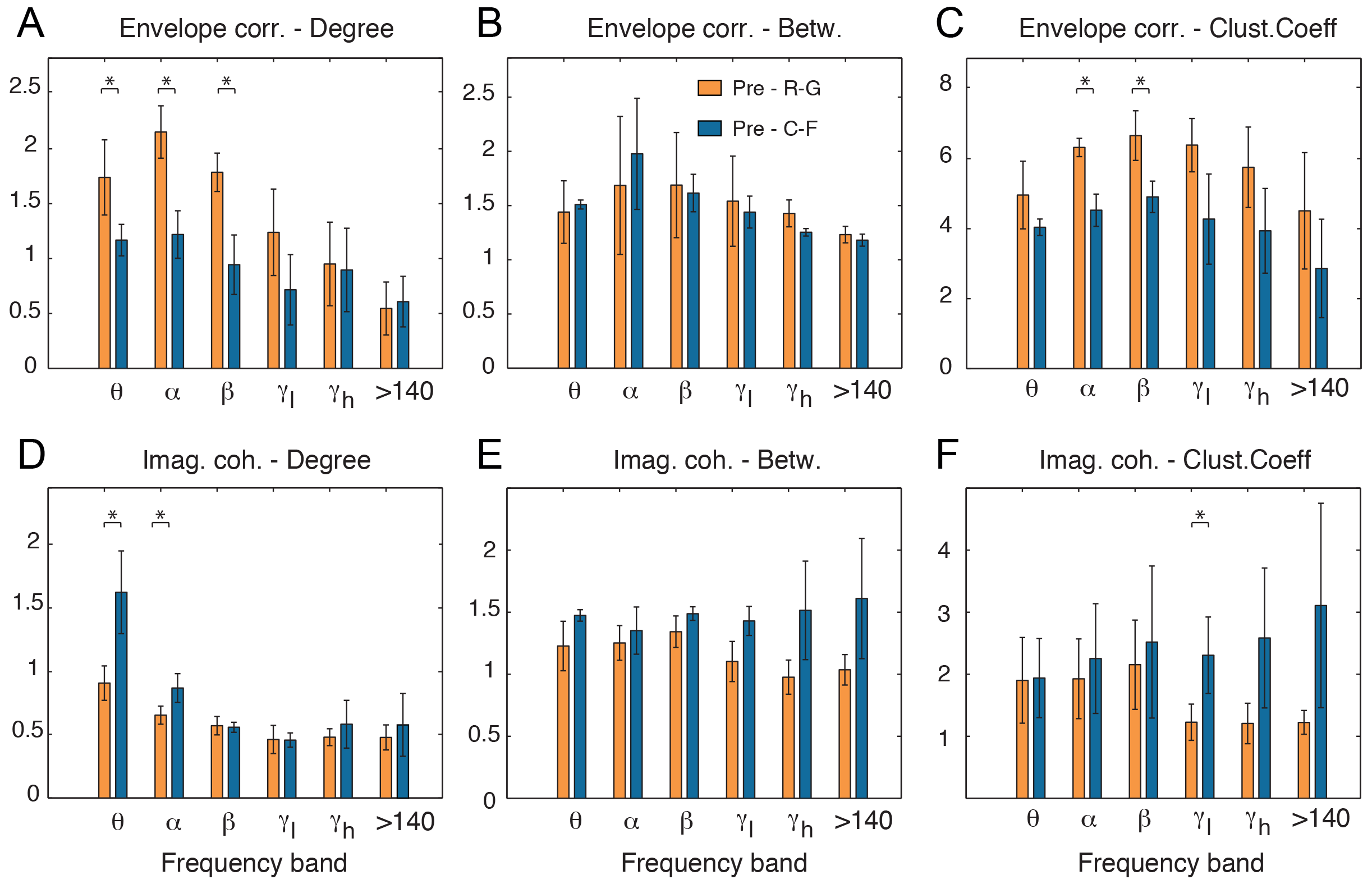
Graph-theoretical analysis of pre-stimulus connectivity. Mean degree, betweenness and clustering coefficient were employed to characterize the ICMs in intervals before stimulus onset. Panels (**A**)-(**C**) show these three measures for envelope ICMs during R-G stimuli (orange) and C-F blocks (blue) across frequency bands. Significant differences occur at low frequencies, in particular theta, alpha and beta bands. Asterisks represent p-values < 0.001. Panels (**D**)-(**F**) show the graph-theoretic measures characterizing the prestimulus intervals in C-F and R-G blocks. The data presented are the mean ± s.d. across animals.

We checked whether these differences in pre-stimulus connectivity were a consequence of differences in the power spectral distribution between pre-stimulation conditions. Fig. S2A displays the mean power distributions of both stimulus conditions across animals and recording sites. Both stimulus conditions showed, on average, very similar distributions with no significant differences across the frequency ranges of interest.

The observations on the pre-stimulus blocks led us to investigate the global structure of connectivity patterns during long periods devoid of stimulation. In addition to the sensory stimulation recordings, in each animal we recorded 3-4 periods of ongoing activity randomly positioned relative to the stimulation blocks. From these periods, which were of at least 15 minutes duration, we sampled 100 randomly distributed epochs of 500 ms duration to match the trial structure during the stimulation blocks. We did not extract epochs during the first 3 minutes of the ongoing activity blocks to minimize any possible effect resulting from preceding stimulation. Comparison of the connectivity matrices for ongoing activity blocks and pre-stimulus intervals during the stimulation blocks yielded a number of observations (Fig. 4). First, envelope ICMs in ongoing activity resembled, in terms of their strength, more closely those in pre-sustained than in pre-transient intervals. Within the auditory areas, the pre-sustained connectivity showed a slight decorrelation relative to the ongoing condition (e.g., sustained mean = 0.08 vs. ongoing mean = 0.150; t-test, *p* < 10^−6^), which extended across all frequency bands (Fig. 4). In contrast, phase ICMs during ongoing activity showed larger similarity in strength to the pre-transient condition at low frequencies. Analysis of the connectivity matrices after normalization to the mean connectivity in each matrix (Fig. S6) revealed that, for envelope ICMs, the spatial pattern of coupling was qualitatively similar across the different blocks for some frequency bands (e.g., the low and high gamma band) whereas for other bands differences were still apparent (e.g., the alpha band).

Taken together, our results indicate that it is highly probable that differences in connectivity between stimulus-blocks are not the consequence of network responses to individual stimuli but the result of a more global change that may relate to the stimulation context on longer time scales. This suggests that during the presentation of a train of stimuli the brain undergoes a context specific reconfiguration of functional networks that can differ from the spatial pattern of coupling modes that prevail during long periods without any sensory stimulation.

### Pre-stimulus ICMs predict multisensory response effects

In a final step of our analysis we investigated the role of these context-specific network reconfiguration. We hypothetized that the multisensory effects observed during stimulus processing, as reflected in the latency and power changes described above, might be a consequence of this functional reconfiguration. For both types of ICMs we tested whether there was a correlation between the strength in the pre-stimulus connectivity and the change in latency (lat_V_ – lat_AV_) observed in visually responsive sites (Fig. 6A,B; cf. Fig. 1E). We included pre-stimulus connectivity between sites that responded only to visual stimulation and sites that responded exclusively to auditory stimuli. For these connections, which involved mostly auditory and early visual areas (Fig. S7A,B), we computed the Spearman-correlation between pre-stimulus connectivity strength and the change in latencies during the subsequent response. We observed that this correlation was specific to both frequency bands and coupling measures (Fig. 6). During the blocks with transient sensory stimulation, beta and low-gamma envelope ICMs induced a reduction of latencies in visual responses (Fig. 6A, *p* < 10^−4^; Pearson’s correlation with Bonferroni correction). Phase ICMs predicted a latency reduction mainly during the sustained stimulation blocks (Fig. 6B). Here, change in response latencies were positively correlated with the pre-stimulus phase-coupling strength in alpha, beta, and low gamma band (*p* < 10^−3^; Pearson’s correlation with Bonferroni correction). The overall comparison of these results across stimulus blocks and coupling measures showed that, except for the correlation in beta band, all significant correlations were either in one or the other coupling mode, suggesting that phase and envelope ICMs may reflect independent coupling mechanisms that potentially differ in the functional relevance.

**Figure 6.**
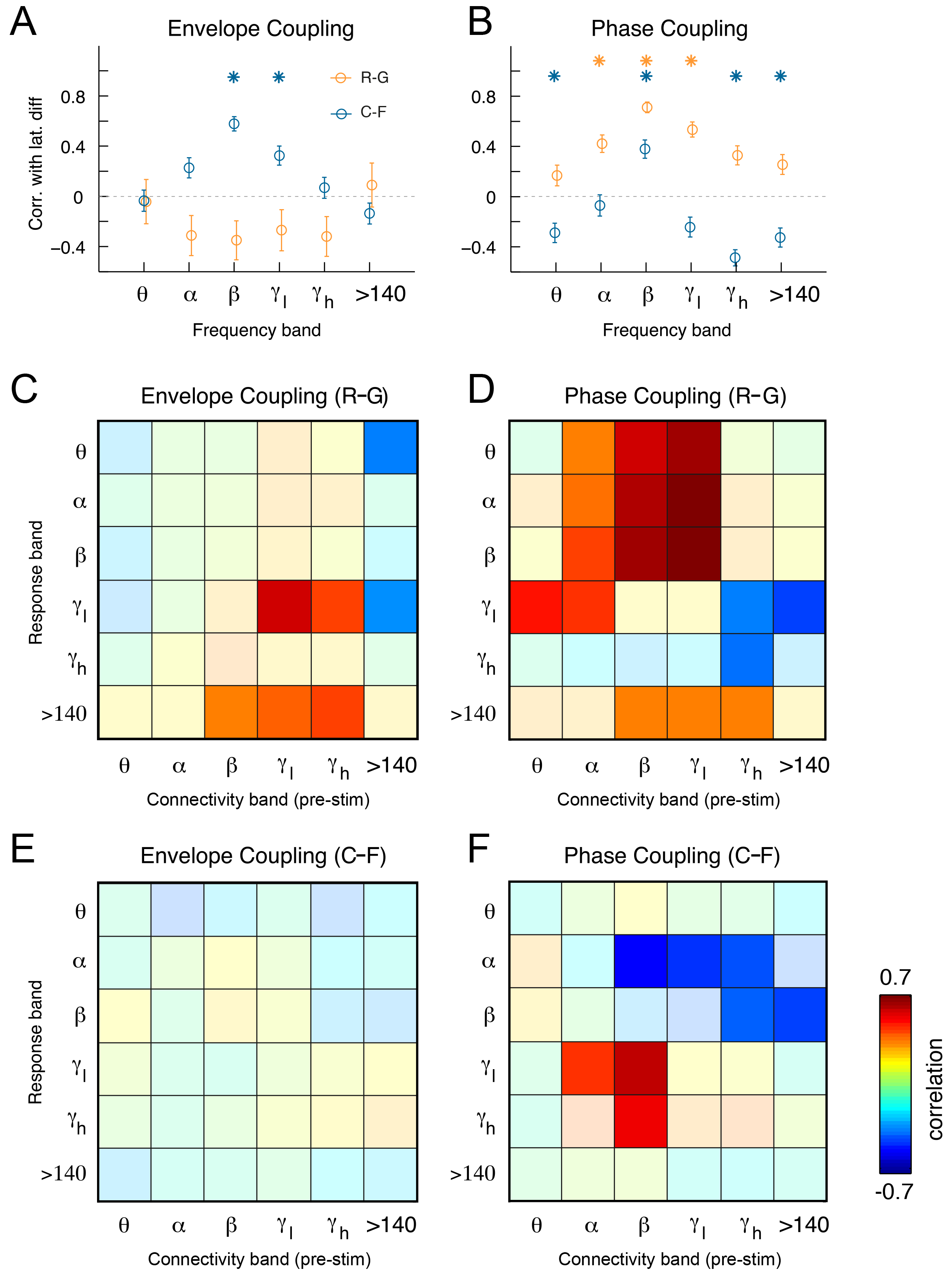
Pre-stimulus connectivity predicts multisensory effects on response timing and power. (**A**) Correlation between envelope coupling and change in latencies (latV-latAV) for visually responsive sites in the R-G (orange) and C-F (blue) stimulation blocks. Asterisks represent significant correlations (p < 0.01), and error bars represent the standard error of the correlation. (**B**) Correlation of differences in latency with phase coupling. (**C**) Correlation between pre-stimulus envelope coupling and multisensory power enhancement observed with ripples and Gabor patches (R-G). Each element in the matrix represents the strength of the correlation between the pre-stimulus connectivity in a certain band, and the stimulus-related power enhancement in a specific band. Hot colors represent positive correlations and cold colors negative correlation. Non-significant correlations (p > 0.01) are masked. (**D**) Correlation matrixs between pre-stimulus phase coupling and stimulus related power changes with R-G. (**E,F**) Same analyses of the relation between pre-stimulus connectivity and multisensory power enhancement for the stimulation blocks with clicks and flashes (C-F).

Following a similar hypothesis, we studied whether the strength of pre-stimulus connectivity predicted multisensory effects on response power. Note that, in contrast to latency, both response power and connectivity can be defined in a spectrally specific manner. Therefore, results of this analysis are displayed as a matrix in which rows represent the response power frequency band and colums represent the pre-stimulus connectivity frequency band (Fig. 6C-F). The results show specific patterns of effects differing across coupling modes, frequency bands and stimulation blocks. In the sustained stimulation blocks, prestimulus envelope ICMs in the low and high gamma band were positively correlated with multisensory responses in the low gamma band (Fig. 6C) (Pearson’s correlation *r* = 0.6, *p* < 0.01, Bonferroni corrected for number of elements of the matrix). In contrast, high-frequency envelope ICMs negatively correlated with multisensory response power changes (Fig. 6C). These effects were not observed in the blocks with transient sensory stimulation (Fig. 6E). However, the strongest correlations occurred for phase ICMs (Fig. 6D,F). In particular, phase ICMs in frequency ranges between 4-64 Hz (theta to low gamma) correlated with multisensory effects at frequencies higher than 30 Hz in the sustained blocks. In the transient blocks a relation was observed between alpha and beta phase ICMs and gamma band response effects (Fig. 6F). Interestingly, substantial negative correlations occurred in the transient stimulation blocks between pre-stimulus phase coupling higher than beta, and multisensory response effects in the alpha and beta ranges (Fig. 6F). It is interesting to note that for response frequencies >140 Hz the correlations with broad band pre-stimulus connectivity were significant for both types of coupling modes in the sustained stimulation blocks. In contrast, transient stimulation blocks showed a very different pattern of correlations (Figs. 6E and 6F), in which a predictive relation for high-frequency (>140 Hz) components was mostly absent.

Taken together, these results suggest a causal effect of pre-stimulus large-scale ICMs on multisensory response power effects, in which functional connectivity between early sensory areas before the stimulus presentation seems to play an important role.

## DISCUSSION

This study has investigated the interplay between intrinsically generated large-scale coupling and stimulus-related activity using an audiovisual paradigm in the ferret. We hypothetized that repetitive sensory stimulation reconfigurates the ICMs, and that this reconfiguration at the same time affects the upcoming sensory stimulus processing. Our approach was to present auditory, visual and audiovisual stimuli to study whether such reconfiguration could act as facilitator during multisensory integration. We tested this hypothesis in an anesthetized preparation, to have a better control on the sensory inputs and to reduce effects of other factors or brain states such as arousal, attention and motor preparation. We used an ECoG array to record brain activity under two different stimulus blocks, and during periods without any sensory stimulation, to study whether this possible reconfiguration of ICMs might also occur spontaneously or is guided by the context.

Analysis of the topography of responses to visual and auditory stimuli showed a power increase in response to unimodal stimuli which spread across several frequency bands, including alpha, which is reduced in response to visual stimuli in awake conditions (39). Multisensory stimulation also produced changes in stimulus-related power that showed both enhancement and mostly suppression across frequency bands. Enhancement was dominant in visual areas (beta and low-gamma), presumably area 20, and at high frequencies (> 64 Hz) in the secondary auditory areas ADF and PPF, which possibly correspond to the borders between unimodal subregions within AES area in the cat (31, 33). It is important to note that, due to the inverse effectiveness of multisensory integration (40), this distribution of suppression/enhancement may have been favoured by the stimuli applied here, which were presented at moderately high intensities to assure the sensory responses (see Methods). Interestingly, the topographic maps of suppression and enhancement appeared to be stimulus block specific (35). In our study the responses to auditory and visual stimuli were not exclusive to their associated functional pathways, but it was frequent to observe visual responses in auditory areas, and vice versa. This crossmodal effect has been widely reported across the cortex in several functional areas and species (30, 35, 41, 42). Finally, because of the relatively short interstimulus intervals, sensory habituation might be expected. However, effects of recent history in the network response were not observed (Fig. S1). The combination of different stimulation modalities as well as the randomized interstimulus intervals may contribute to a decreased efficiency of habituation mechanisms.

A recurrent result was that latencies at visually responsive cortical sites were shortened by combined audiovisual stimulation compared to the unimodal visual condition. This was observed, in particular, for higher-order sensory regions around the suprasylvian sulcus and for regions around the pseudosylvian sulcus. We interpret this as the consequence of a collaborative interaction between both modalities. This reduction in latencies might be responsible for the speeding up of perceptual and motor responses to visual targets (43–47). In our study, the presence of the visual stimuli did not affect the latency of auditory responses, as has been observed in awake monkeys (48). One possible reason for this may be that our recordings were done in the anesthetized preparation. Another reason why shortening in auditory responses were not observed may be related to the fact that visual and auditory stimuli were presented in synchrony. If occurring asynchronously, visual stimulation can reset the phase of ongoing auditory cortical oscillations, and the degree of asynchrony can regulate the enhancement or suppression of multisensory responses (35, 49).

Oscillatory dynamics and frequency-specific coupling across multiple temporal scales are important characteristics of functional networks in the brain. Two of the mechanisms that underlie intrinsic coupling modes are phase coupling of band-limited oscillatory signals (phase ICMs) and the correlated fluctuations of signal envelopes (envelope ICMs) (1). Our results demonstrate that envelope ICMs and phase ICMs differ strongly in their spatial patterning. Whereas envelope ICMs showed a clustering of areas into functional systems, phase ICMs tended to be higher for connections between, rather than within functional clusters of cortical areas. For envelope ICMs, the spatial topography of coupling patterns seemed to be similar across frequency bands, but the strength of coupling varied across frequency ranges. The mean correlation values of envelope ICMs were higher in theta, alpha and beta bands than in higher frequency bands. Therefore, we observed a decay of average connectivity as frequency increases and, at least in blocks with sustained stimulation, this was significantly lower at frequencies above 140 Hz. Although not in such a drastic manner, the same result was observed in the transient stimulation blocks. This agrees with the idea that envelope coupling captures mostly correlated fluctuations at lower frequencies, in a manner similar to what has been observed by correlation of fMRI BOLD signals (50, 51). This dependence of coupling strength on frequency bands was much less prominent for phase ICMs. This fact, together with the strong difference in patterns across the two coupling modes supports the idea that these types of ICMs may mediate different aspects of the dynamic communication across brain areas. Whether they mutually interact or are independent needs to be further investigated.

To our surprise, the stimulus-related connectivity patterns did not appear to depend on stimulus modality, and they did not differ substantially from the pre-stimulus interval for any of the modalities and coupling modes. Interestingly, comparison of connectivity matrices between stimulation blocks during the pre-stimulus windows revealed prominent differences suggesting that, rather than the current stimulus itself, it was the *global context*, created by the prolonged repetition of similar stimuli, that strongly shaped ICMs. This suggests that the long-term stimulation context can induce specific network states reflected in the dynamics of large-scale coupling patterns. Comparison of the spatial patterns for envelope and phase ICMs revealed a clear dissociation of the two types of coupling modes. This suggests that network connectivity states can emerge independently through the coordinated coactivation of sensory areas on slower time scales (as reflected in envelope ICMs) or through phase alignment of neural signals (as reflected in phase ICMs). Importantly, this was observed only with respect to functional connectivity and not regarding the local spectral properties of ongoing activity which did not show significant differences across stimulation contexts. Our findings suggest a novel form of brain state reconfiguration that, in contrast to previous studies (12, 27), affects connectivity without altering the energy of local signals. This reconfiguration seems to occur as a result of prolonged exposure to certain sets of sensory stimuli, leading to a reshaping of the predominant patterns of both envelope and phase ICMs.

Such a context-dependent reconfiguration of functional networks, in turn, might have an impact on the processing of individual sensory stimuli. Previous studies have shown the impact of pre-stimulus states on multisensory interactions (13, 49, 52). We tested this by correlating envelope and phase ICMs between visual and auditory regions in pre-stimulus epochs with the multisensory effects that we observed on latency and power of local responses. This analysis revealed strong effects of pre-stimulus functional connectivity on multisensory stimulus processing. Interestingly, the impact of pre-stimulus connectivity on multisensory responses differed between ICM types and stimulus contexts. Thus, for instance, response latency reductions were predicted mainly by envelope ICMs in the transient stimulation blocks, but by phase ICMs in the blocks with sustained stimulation. Multisensory effects on response power were predicted mainly by phase ICMs. The ICMs predicting timing and power changes during multisensory stimulation predominantly involved connections between primary auditory and visual areas, supporting the notion that reconfiguration of connectivity as a mechanism for multisensory integration may already take place at early cortical processing stages (53).

Our study suggests that long-term modulation by repetitive sensory inputs can lead to a context-specific shaping of intrinsic coupling modes. These reconfigured coupling modes, in turn, lead to changes in latencies and power of local field potential responses that support multisensory integration. Our results demonstrate that this reciprocal interplay between network dynamics and external signals extends across multiple frequency bands and involves different types of intrinsic coupling. Although the present study was done in an anesthetized preparation, we suggest this large-scale contextual adaptation may provide a mechanism which effectively facilitates responses of the network to sensory stimulation, as observed previously in behavioral studies (54). This might be particularly suitable for processing of complex stimuli which require adaptation to the statistics of the environmental signals. In conclusion, the present study sheds light on the relationships between large-scale ICMs on multiple time scales and cortical evoked responses, and provides evidence for the basis of a large-scale facilitating mechanism for multisensory integration of complex stimuli.

## MATERIALS AND METHODS

Data presented in this study were collected from five adult female ferrets *(Mustela putorius)*. All experiments were approved by the Hamburg state authority for animal welfare (BUG-Hamburg) and were performed in accordance with the guidelines of the German Animal Protection Law.

### Surgery

Animals were initially anesthetized with an injection of ketamine (15 mg/kg) and medetomidine (0.08 mg/kg). A glass tube was then placed in the trachea to allow artificial ventilation of the animal and to supply isoflurane anesthesia (0.4% isoflurane, 50:50 N_2_O – O_2_ mix). To prevent dehydration throughout experiments, a cannula was inserted into the femoral vein to deliver a continuous infusion of 0.9% NaCl, 0.5 % NaHCO_3_ and pancuronium bromide (60 μg/kg/h). Body temperature was maintained at 38 °C with a heating blanket controlled by measurement of the animal’s rectal temperature. Heart rate and end-tidal CO_2_ concentration were constantly monitored throughout the duration of experiments to maintain the state of the animal. Before fixing the animal’s head in the sterotaxic frame, a local anesthetic (Lidocain, 10%) was applied to the ear bars. To keep the animal’s head in a stable position troughout the placement of the ECoG and the measurements, and allow auditory and visual stimulation, a headpost was fixed with screws and dental acrylic to the frontal bone of the head. The temporalis muscle was folded back, so that a large craniotomy could be performed over the left posterior cortex. After careful removal of the dura, the cortex was covered with saline solution. The ECoG array was gently placed on the surface of the cortex, such that electrodes covered large areas of visual and auditory cortex (Fig. S1). A small piece of artificial dura (Lyoplant; B. Braun Melsungen AG, Melsungen, Germany) was cut in the shape of the craniotomy and gently placed over the ECoG array. Subsequently, the removed piece of scull was placed over the artificial dura, and then the craniotomy was resealed using silicon elastomer (World Precision Instruments, Sarasota, FL). To prevent exsiccation of the cornea, a contact lens was placed over the right eye, while the left eye was occluded to ensure monocular stimulation. Experiments typically lasted between 24 hours and 36 hours, after which the animal was deeply anesthetized with 5% isoflurane and sacrificed with an overdose of potassium chloride.

### Electrophysiology

During electrophysiological recordings isoflurane level was maintained at 0. 4%. Neural activity across the entire posterior cortex was recorded using a custom designed micro electrocorticogram array (ECoG) (55). ECoG arrays were polyimide based, and consisted of 64 hexagonally spaced (1.5 mm inter-electrode distance) platinum electrodes with a diameter of 250 μm (Fig. S1). Cortical surface LFPs were referenced to a clamp placed on the deflected temporalis muscle. Signals were digitized at 1395.1 Hz (0.1 Hz high pass and 357 Hz low pass filters, respectively), and sampled with an AlphaLab SnR^TM^ recording system (Alpha Omega Engineering, Nazareth, Israel). All analyses of neural data presented in this study were performed offline after the completion of experiments. To obtain the location of ECoG electrodes for each animal, the position of the ECoG on cortex was recorded by taking photographs with a Zeiss OPMI pico microscope. The position of ECoG grids was then projected onto a scaled illustration of a model ferret brain that contained a map of the functional specialization of all posterior cortical areas (20 in total) (30). Data from each electrode was then allocated to the cortical area directly underlying the corresponding ECoG contact. To avoid potential confounding effects of flucutations in brain state we kept the anesthesia level as constant as possible and presented the stimulation blocks in a randomized manner.

### Sensory stimulation

To ensure controlled conditions for sensory stimulation, all experiments were carried out in a dark sound-attenuated anechoic chamber (Acoustair, Moerkapelle, Netherlands). Auditory and visual stimuli were generated using the Psychophysics Toolbox (56) in Matlab (Mathworks Inc, Natick, MA). Stimuli were digitalized at 96kHz and delivered with an RME soundcard (RME HDSPe AIO Intelligent Audio Solutions) through a Beyerdynamic T1 speaker located 15 cm from the animal’s right ear. Before performing any experiments, the sound delivery system was calibrated using a Brüel and Kjær (B&K, Nærum, Denmark) free-field 4939 microphone coupled to a B&K 2670 preamp and 2690 amplifier. Visual stimuli were presented on an LCD monitor (Samsung SyncMaster 2233, frame rate 100 Hz) placed 28 cm in front of the animal.

We presented two types of stimulation blocks that strongly differed in the duration and structure of the presented stimuli: (1) A *transient stimulation block* comprised visual stimuli consisting of 10 ms flashes of a 14°x14° white square located at the center of the monitor displayed on a black background, as well as auditory stimuli consisting of 0.5 ms sound clicks presented at 65 dB SPL. We presented both, unimodal and bimodal conditions (with zero delay) in a randomized intermixed fashion with inter-stimulus intervals varying between 900 and 1100 ms. In total, 100 flash, 100 click and 100 simultaneous clicks and flashes stimuli were presented per stimulation block. (2) A *sustained stimulation block* comprised drifting Gabor patches with a size of 14 degrees (full width at half maximum), a spatial frequency of 2 cycles/deg and a temporal frequency of 5 deg/sec, drifting in 4 directions (0, 90, 180 and 270 degrees). For the auditory stimulation we presented dynamic moving ripples, an acoustic analog of the visual gratings (57). Moving ripples are complex noise-like stimuli that consist of the sum of several sinusoidal amplitude-modulated tones. By adjusting amplitude and phase of the tones they obtain a sinusoidal spectral envelope (58). We created the ripples using 60 tones logarithmically spaced along the frequency axis between 0.5 – 32 kHz. The spectral envelope of the composite sound was then modulated as a single sinusoid along the frequency axis on a linear amplitude scale (spectral peak density Ω = 0.1 and 1 cycle/octave; and peak drift speed w = 0.5 and 0.2 Hz in both directions). Acoustic signals were multiplied by a 0.5-ms cos^2^ onset and offset function, and scaled to a target root-mean-square (RMS) amplitude to generate and present stimuli at 65 dB SPL. In total, 100 Gabor patches, 100 ripple stimuli and 100 simultaneous patches and ripples were presented per stimulation block.

Stimuli of the sustained blocks had a duration of 1000 ms with interstimulus intervals varying between 900 and 1100 ms. Finally, to enable comparison of results obtained from pre-stimulation intervals during the two types of stimulus blocks with longer epochs not perturbed by sensory stimulation, we also recorded blocks of 10 to 15 minutes duration of ongoing activity. We recorded 3 to 5 of these periods per animal at the beginning of recording session and between blocks of stimulation.

### Data analysis

All data analysis was performed using custom scripts in Matlab (Mathworks). Artificial ventilation and heart activity introduce in some cases strong artifacts. For that reason frequencies below 4 Hz were not considered for current analysis. Sensory response latencies were defined as the post-stimulus time where LFPs exceeded four times the standard deviation of pre-stimulus activity. The amplitude of responses was quantified as the largest LFP deflection (positive or negative) in the first 80ms post-stimulus. Spectral components of neural activity following sensory stimulation were estimated by computing fast Fourier transforms (FFT) on LFPs. FFTs were computed from 100-600ms post-stimulus to ensure that transient onset responses did not confound spectral estimates. Reflecting the natural segregation of cortical rhythms into distinct frequency ranges, spectral estimates were averaged within the theta (4-8 Hz), alpha (8-16 Hz), beta (16-32 Hz), low gamma (32-64 Hz), high gamma (64-128 Hz) and high frequency band (>140 Hz). FFT-derived spectral estimates were then used for computing total power of local responses. To this end, we normalized the stimulus-related total power to the pre-stimulus power, which was calculated between −600 and −100 ms relative to stimulus onset. To quantify multisensory effects on neural responses, we computed the raw differences between latencies and also between power of responses to unimodal and bimodal stimulation. For graphical representation of this effect (Fig. 1E,F), we used a variation index which corresponds to the raw difference multiplied by 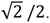

Spectral coherence is a measure of the consistency of phase coupling between simultaneously recorded signals and comprises both real and imaginary components. We chose to quantify phase ICMs by taking only the imaginary part of coherence. This approach eliminates the contribution of volume-conducted signals to functional connectivity measures by exclusively considering non-zero phase lagged signal components (38).

For analysis of envelope ICMs, amplitude envelopes were computed in a time-resolved fashion by band-pass filtering and then Hilbert transforming LFP signals. To avoid the detection of spurious envelope correlation due to the effects of shared noise or volume conduction, pairs of ECoG signals were orthogonalized prior to the computation of the amplitude envelope time series (36). Similar to imaginary coherence, this step removes zero-phase lagged components that are shared between simultaneously recorded signals, therefore ensuring that only non-zero phase-lagged signal components are considered for analysis. Following orthogonalization, amplitude envelopes for the time-window 100-600 ms post-stimulus for all trials were computed. For each stimulus condition, amplitudes were concatenated across all trials. Amplitude correlation was then defined as the linear correlation coefficient of pairwise envelope time series.

Functional connectivity measures were grouped into matrices where each pixel represented the mean functional connectivity for all pairs of electrodes between specialized cortical areas (see below for functional cortical area parcellation). Graph theoretical analysis of connectivity matrices for phase and envelope ICMs was performed using the Brain Connectivity Toolbox (37). To quantify large-scale cortico-cortical connectivity, we computed average degree, betweenness centrality, and clustering coefficients of weighted functional connectivity matrices. The standard deviation of graph theoretical measures was estimated by repeating analysis on randomly shuffled functional connectivity matrices (100 repetitions). Graph theoretical measures were then normalized by the mean and standard deviation of randomly shuffled matrices.

We calculated imaginary coherence and amplitude envelope correlation first on an electrode-to-electrode basis, resulting in a 64×64 matrix with all possible connections. We assigned to each electrode the area over it was located, and then regrouped and averaged connectivity data in terms of pairs of areas, yielding a connectivity matrix in terms of a functional area representation (Fig. 1A). Note that in this new representation the number of electrodes per area varied across areas (from 2 to 6 electrodes). We took averages across animals only for areas that were recorded in all animals. Note that areas V1, AVF and VP, displayed in Figure 1, are not in the mean matrix, because they were not recorded in all animals.

Finally, we evaluated the potencial artifactual connectivity introduced by the underlying volume conductivity of the tissue. To this end we calculated both connectivity measures in two preprocessing conditions: first, using the filtered signals without re-referencing, and second, using local bipolar derivatives (re-referencing to closest neighbouring electrode). Our assumption was that the volume conductivity increases connectivity and, as a consequence, the preprocessing condition less affected by volume conduction shows lower values in both connectivity measures. We measured mean connectivity between all first, second and third neighbouring pairs (Fig. S8). Based on this analysis in the current study we used unreferenced signals.

## ACKNOWLEDGMENTS

We would like to thank Dorrit Bystron for her assistance with data acquisition, and Christian Moll and Alessandro Gulberti for discussions and critical comments.

## Author contributions

EGL, AKE, IS, FP and GE conceived and designed experiments. EGL, IS and FP performed experiments. EGL analyzed data. GE wrote the animal ethics permission. TS designed and developed ECoG arrays. EGL and AKE wrote the paper. All authors edited and approved the manuscript.

## Data and materials availability

All data needed to evaluate the conclusions in the paper are present in the paper and/or the Supplementary Materials. Additional data related to this paper may be requested from the authors.

