## Supplemental information for "Context-specific modulation of intrinsic coupling modes shapes multisensory processing"

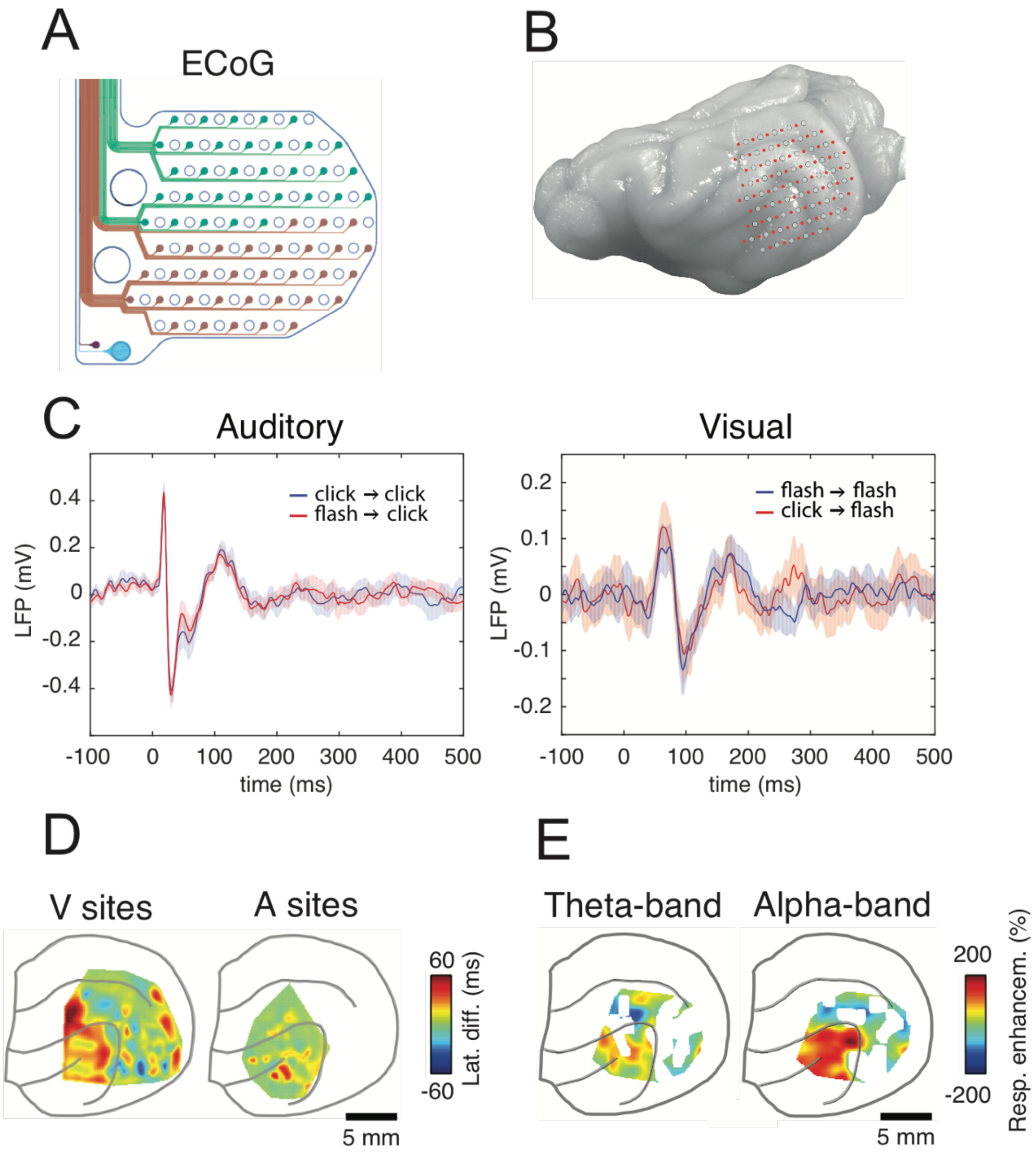


**Figure S1. Recording approach in the anesthetized ferret.** (**A**) ECoG array developed for the current study. A total of 64 platinum thin-film electrodes were integrated into a 10 µm thin polyimide foil. The diameter of each electrode was ~0.25 mm, and the separation between electrodes was 1.5 mm. (**B**) Schematic of the ECoG array placed on the left hemisphere of the ferret brain, covering most of the occipital, temporal and parietal cortex. Red dots represent recording sites, and white circles represent the holes in the foil. (**C**) Effect of preceding stimulus on current response. *Left*: example of responses to clicks in auditory areas in trials depending on whether the preceding stimulus was a click (blue) or a flash (red). *Right*: same analysis in a visual area. No evident habituation was observed with stimulus repetition in our stimulus paradigm. (**D**) Topographic distribution of the difference in latencies between unimodal and bimodal stimuli for visually (left) and auditory (right) responsive sites. Electrodes with no response are not included in the topographic maps. (**E**) Topographic distribution of the relative differences in power during R-G stimulation: ($LFP_{AV}$-$LFP_{A}$-$LFP_{V}$), with LFP being the power of the response.

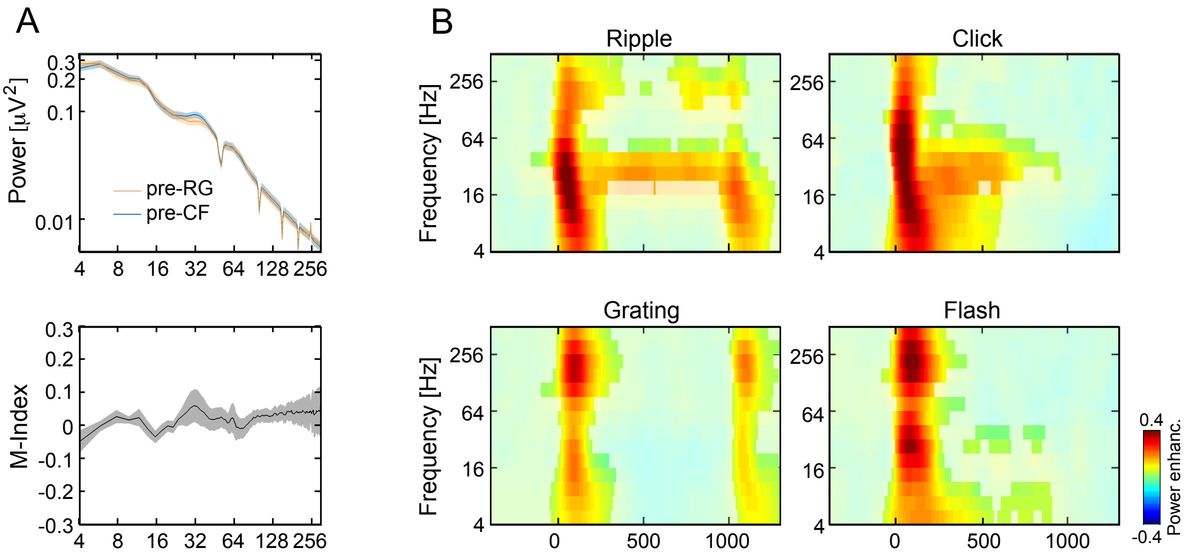


**Figure S2. Spectral properties of ongoing and stimulus-related activity.** (**A**) Upper panel: power spectral density of LFP signals during pre-stimulus windows in blocks of R-G stimulation (orange) and blocks of clicks/flashes (blue). The panel displays the mean power distributions (± s.e.m.). Lower panel: difference between pre-stimulus power between R-G and C-F stimulus blocks, expressed as modulation (M-) index $(LFP_{RG}-LFP_{CF})/(LFP_{RG}+LFP_{CF})$. (**B**) Spectrograms of stimulus-related total power for all unimodal stimuli presented in this study. Spectrograms show the average across sites that responded to auditory stimuli (top row: ripples and clicks) and sites that responded to visual stimulation (bottom row: drifting Gabor patches and flashes). Responses are normalized to the pre-stimulus interval (-600 to -100 ms). Non-significant power changes were masked.

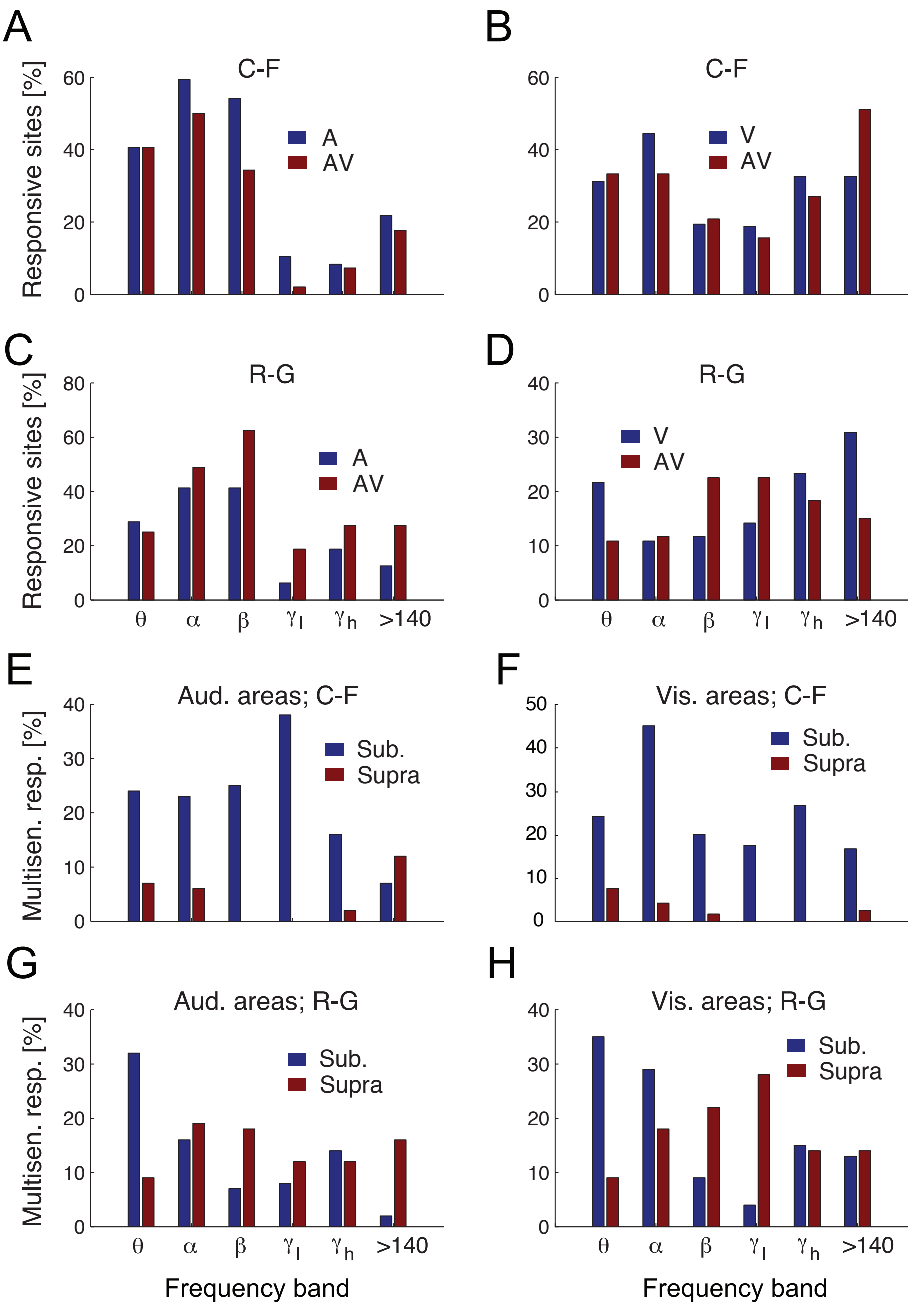


**Figure S3. Fraction of responsive sites and multisensory responses**. (**A**) - (**D**) Proportion of responsive sites (see Methods) in auditory and visual areas for unimodal and bimodal stimuli across frequency bands. (**A** and **B**, clicks-flashes; **C** and **D**, ripples-Gabor patches). We found across the population that the percentage of responsive sites in both sensory modalities was not stimulus dependent and it was not different across stimulus blocks (*P*> 0.05; ANOVA, after Bonferroni correction). However, the number of responsive sites was frequency dependent in auditory areas (*P* = 0.003; n-way ANOVA, Bonferroni corrected), but not in visual areas (*P* = 0.28; ANOVA). (**E**)-(**H**) Proportion of multisensory sites, with multisensory effects divided into superadditive and subadditive response groups for C-F blocks (**E**, **F**) and R-G blocks (**G**, **H**). Across visual and auditory areas, both types of multisensory effects were observed. The subadditive effects were dominant in both sensory systems, in particular at low frequencies. Interestingly, the percentage of sites that showed supraadditive multisensory effects was not frequency dependent (*P* > 0.4 in both cases).

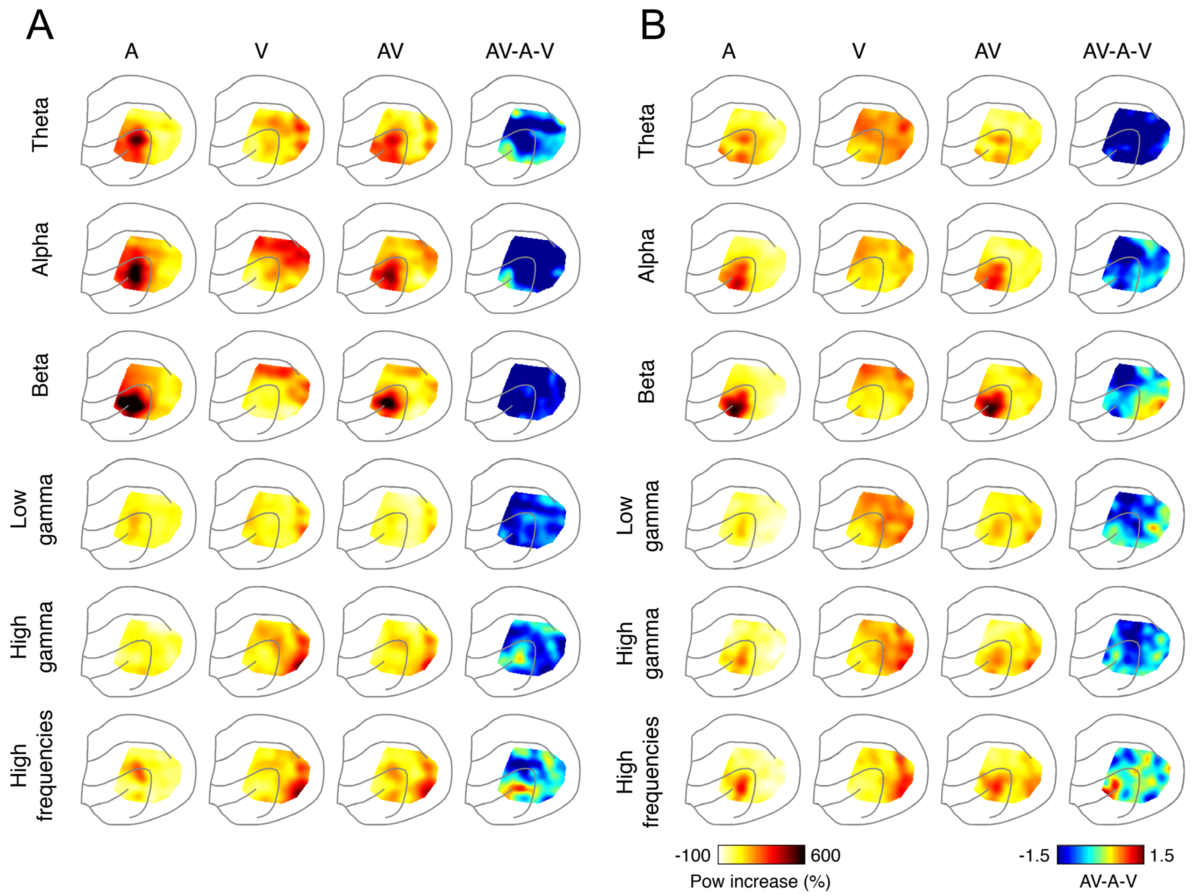


**Figure S4. Topography of response power changes for sustained and transient stimuli**. (**A**) The first to third column show the mean power distribution for unimodal ripples (first column), unimodal drifting Gabors (second column) and the bimodal stimulus (third column). Increase of total power was derived as $LFP_{inc} = (LFP_{st} - LFP_{pre})/LFP_{pre}$, with $LFP_{st}$ and $LFP_{pre}$ being the average power between 100 and 600 ms (relative to stimulus onset) and -600 to -100 ms, respectively. The fourth column displays the mean difference between bimodal power increase and the sum of the unimodal responses. The multisensory effects, defined as the difference in power AV-A-V were calculated first for each animal and then averaged across animals. Regions where this difference was not significantly higher or lower than zero were masked. Rows represent the frequency bands as defined in Methods. (**B**) The first to third column show the mean power distribution for unimodal clicks (first column), unimodal flashes (second column) and the bimodal stimulus (third column). The fourth column displays the mean difference between bimodal power increase and the sum of the unimodal responses.

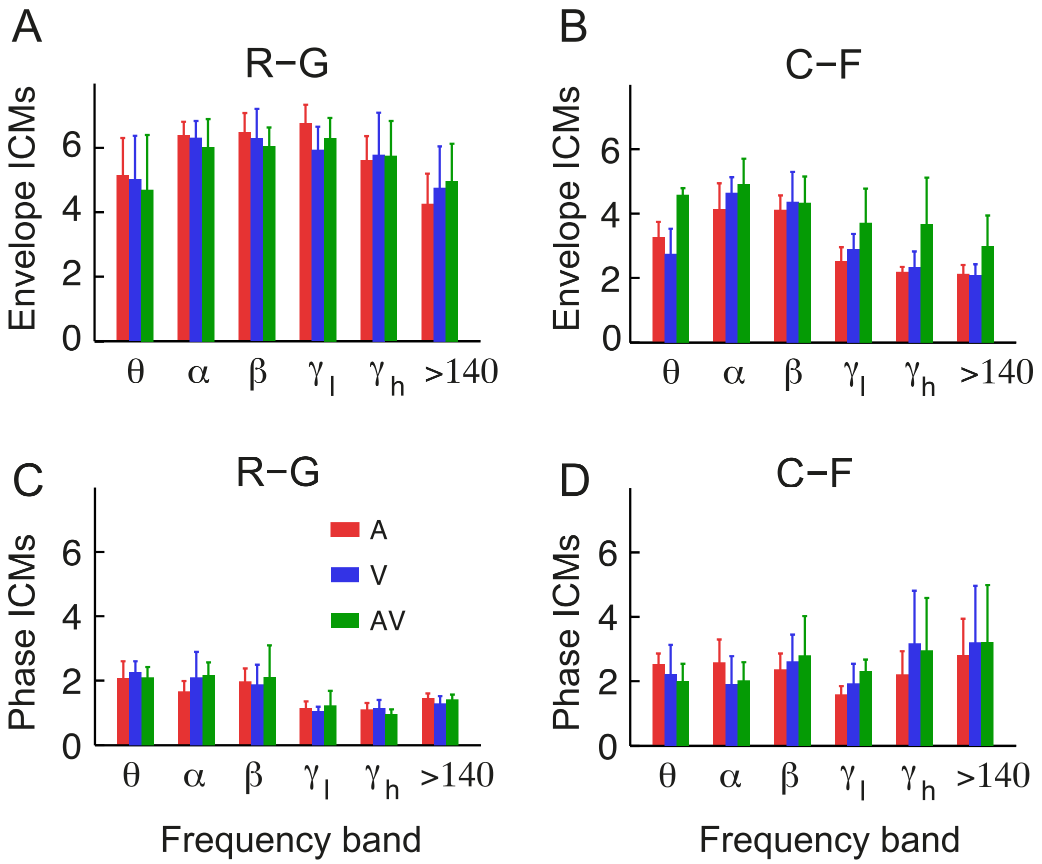


**Figure S5. Clustering coefficient for stimulus-related connectivity.** (**A**,**B**) Results for envelope ICMs in ripple-Gabor (R-G) blocks and click-flash (C-F) blocks. (**C**,**D**) Same analysis for phase ICMs.

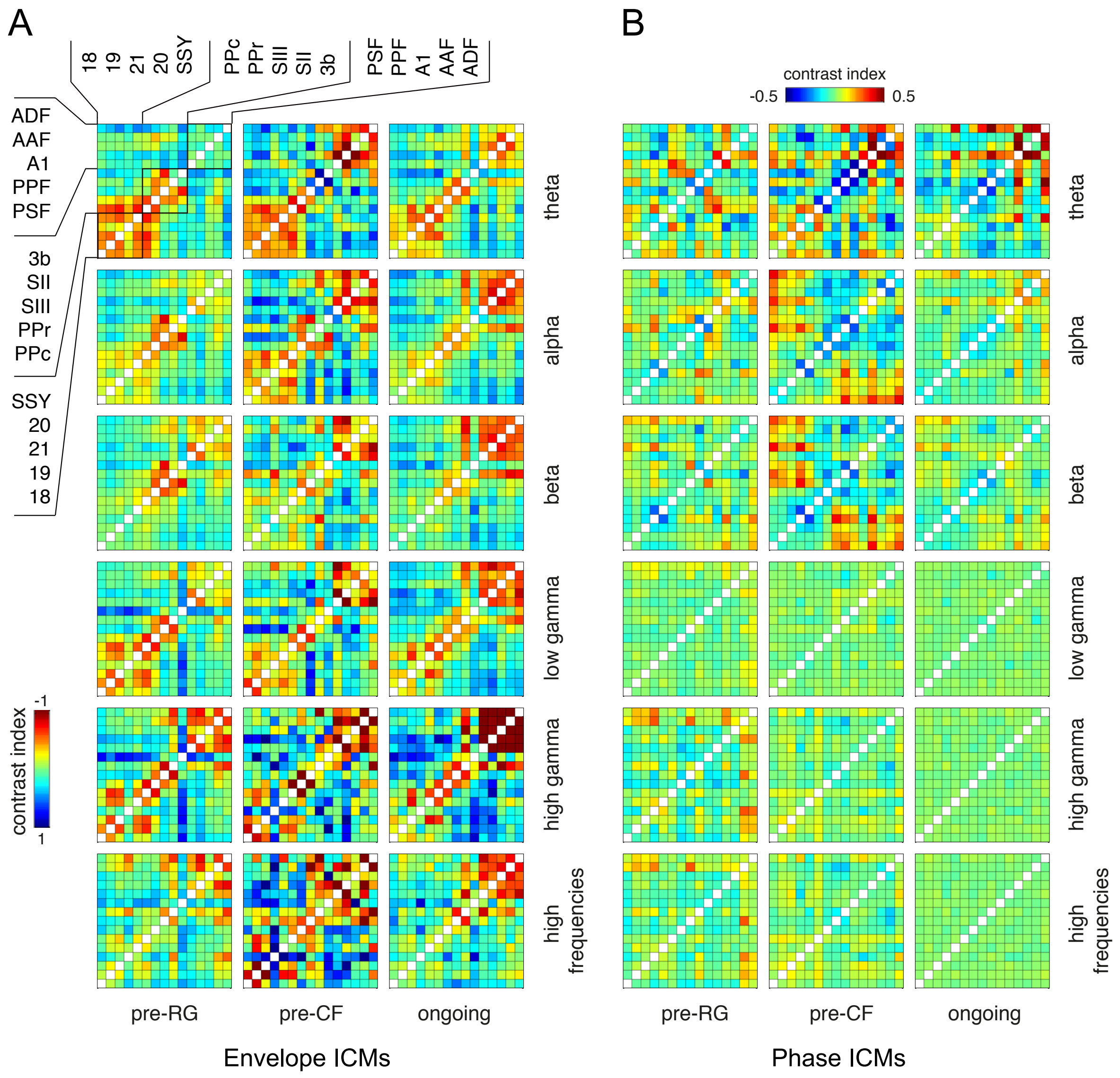


**Figure S6. Contrast index of connectivity matrices during pre-stimulus conditions.** To reveal connectivity positively or negatively exceeding the mean connectivity of the matrix, a contrast (C-) index was defined as M/<M>–1, with M representing the connectivity matrix, and <M> the mean value of the elements excluding the diagonal. (**A**) Amplitude envelope correlation. (**B**) Imaginary coherence. Conventions for rows and columns as in Fig. 4.

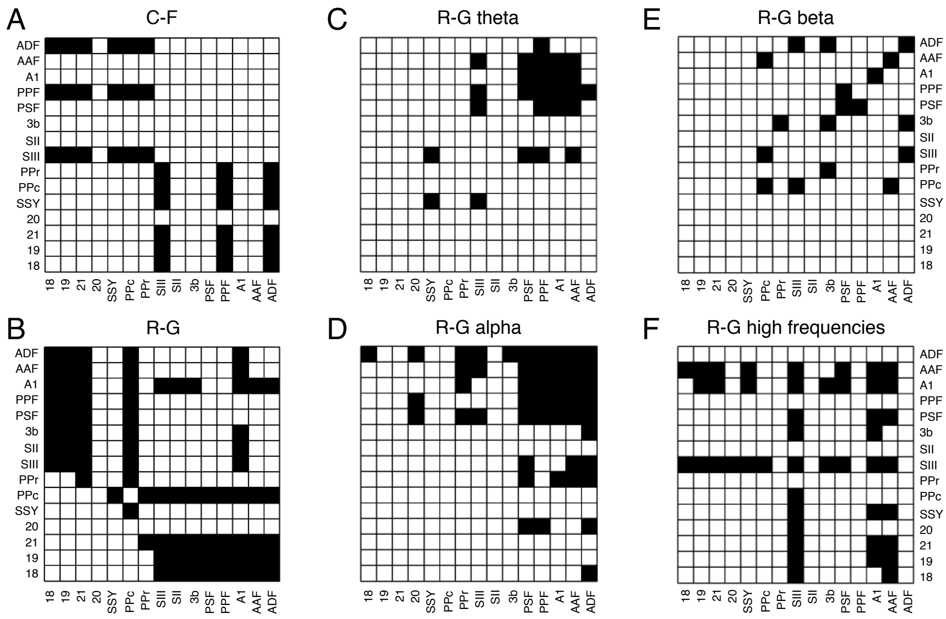


**Figure S7. Relation between pre-stimulus functional connectivity and multisensory effects.** (**A,B**) Binary connectivity matrices showing connections (black bins) employed for the connectivity-latency correlation displayed in Figures 6A and 6B, respectively. (**C**,**D**) Connections involved in the correlation between pre-stimulus phase coupling and multisensory power enhancement in theta and alpha frequency bands. (**E**,**F**) Connections responsible for the correlation between pre-stimulus envelope coupling and multisensory power enhancement in beta and high-frequency bands, respectively.

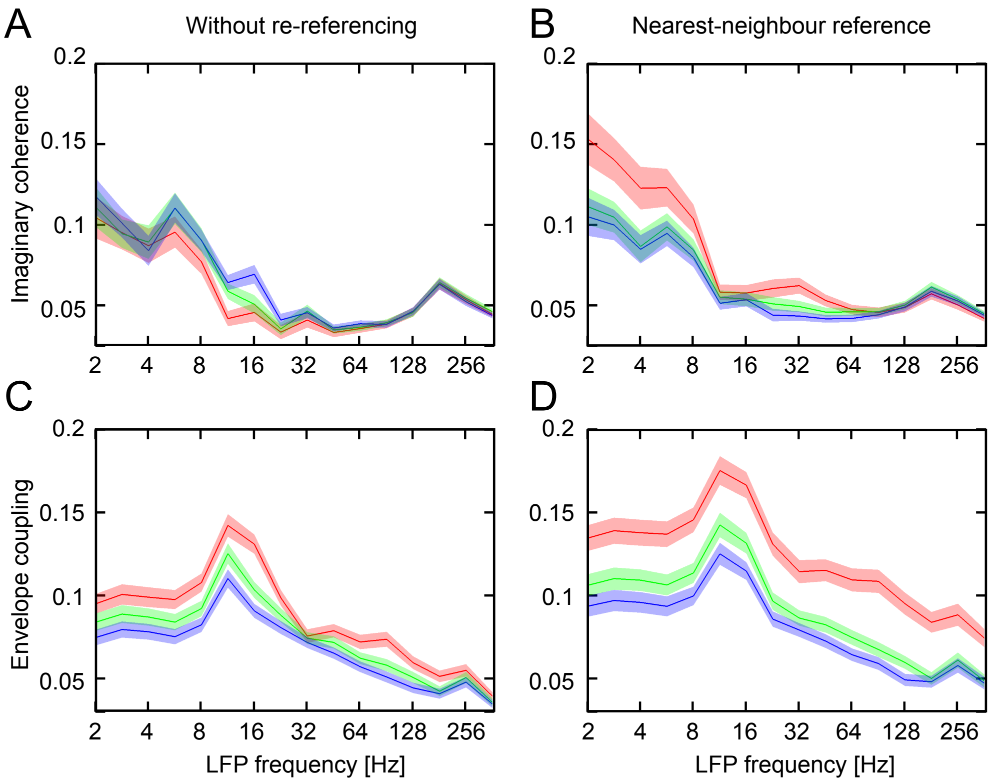


**Figure S8. Grand mean of connectivity measures as a function of frequency.** (**A**) Imaginary part of the coherence for non-re-referenced ECoG signals (raw signal). Red: Mean connectivity between all pairs of first-neighbours (d = 150 µm); green and blue: connectivity between pairs of second (d = 300 µm) and third neighbours (d = 450 µm), respectively. Shading represents the s.e.m. (**B**) Imaginary part of the coherence for re-referenced ECoG signal (nearest neighbour) as function of frequency. (**C**) and (**D**) Amplitude envelope correlation for ECoG signals without and with re-referencing, respectively.
